# Innate potential of random genetic oligomer pools for recombination

**DOI:** 10.1101/320499

**Authors:** Hannes Mutschler, Alexander I. Taylor, Alice Lightowlers, Gillian Houlihan, Mikhail Abramov, Piet Herdewijn, Philipp Holliger

## Abstract

The spontaneous emergence of function from prebiotic pools of informational polymers is a central conjecture of current origin of life scenarios. However, the innate functional capacity of random genetic polymer pools is unknown. Here, we have examined the *ab initio* activity of random and semi-random eicosamer pools of RNA, DNA and the unnatural genetic polymers ANA (arabino-), HNA (hexitol-) and AtNA (altritol-nucleic acids) with respect to a simple functional test: the capacity for intermolecular ligation and recombination. While DNA, ANA and HNA pools proved inert, naïve RNA and AtNA pools displayed diverse modes of intermolecular recombination in eutectic ice phases. Recombination appears linked to the vicinal ring cis-diol shared by RNA and AtNA. Thus, the chemical configuration that renders both susceptible to hydrolysis also enables substantial spontaneous intrapool recombination in the absence of activation chemistry with a concomitant increase in the compositional and structural complexity of recombined pools.

---

**One Sentence Summary:** A vicinal cis-diol configuration, which renders RNA susceptible to hydrolysis, enables spontaneous recombination within naïve random sequence RNA pools.

The phenotypic potential, i.e. the density of functional sequences within oligomeric sequence space, is a central (if largely unexplored) parameter of any given chemistry. This critically informs both the adaptability of biological systems dependent on combinatorial diversity (such as the mammalian immune system) as well as the proposed spontaneous emergence of function from random oligomer pools at the origin of life. The sequence space of RNA, DNA and a variety of xeno nucleic acids (XNAs) contain functional molecules, ligands (aptamers) and catalysts, which can be discovered through iterative rounds of enrichment and amplification. Extrapolation from such selection experiments and analysis of the mutational landscapes of functional oligomers (*1*) suggests that the frequency of e.g. distinct ligands and catalysts is rare (10^−11^ – 10^−14^). However, a focus on defined “hit” sequences may underestimate the global functional potential of oligomer pools, as weak catalysts may be considerably more frequent (*2–4*). The functional potential of a pool may also depend on system-level interactions among pool members such as potentially mutualistic (or antagonistic) interactions, which boost (or interfere) with function (*5*). Finally, function may arise not only from folding of a single oligomer, but also by non-covalent association of short oligomers into an active complex (as observed e.g. for the hairpin ribozyme (HPz) in eutectic ice phases (*6, 7*)).

Here, we have sought to examine the potential of pools of short oligomers of semi- and fully-random sequence of different genetic polymers (RNA, DNA, ANA (arabino-), HNA (hexitol-) and AtNA (altritol-nucleic acids) with regards to a simple test of function: the ability to undergo intermolecular ligation (or recombination) both in the presence and absence of chemical activation, analyzing reacted pools directly by deep sequencing.

## Results

We first examined the reactivity of 2’, 3’-cyclic phosphate (>p)-activated random RNA pools. >p groups form as part of RNA degradation reactions as well as during proposed prebiotic nucleotide synthesis (*8–11*) and mediate non-enzymatic ligation in the presence of an organizing template (*12, 13*). In order to maximize reactivity, we incubated pools in ice, where eutectic phase formation has been shown to promote molecular concentration and interaction, reduce hydrolysis, and provide a compartmentalized medium (*14, 15, 7*). We first incubated a >p-activated random RNA eicosamer pool (N_20_>p) in ice without any organizing template and searched for higher molecular weight (MW) product bands in electrophoretograms from denaturing gel electrophoresis (Urea-PAGE) (Fig 1A). We observed intermolecular covalent N_20_>p pool ligation yields of ~10% (after 2 months, -9 °C) in either presence or absence of MgCl_2_ (Table S3). No ligation was observed in equivalent control samples incubated at -80°C (Fig. 1B, C). Deep sequencing of the prominent higher MW band confirmed that this corresponded to ligation events (Fig. 1C, D).

**Fig 1.**
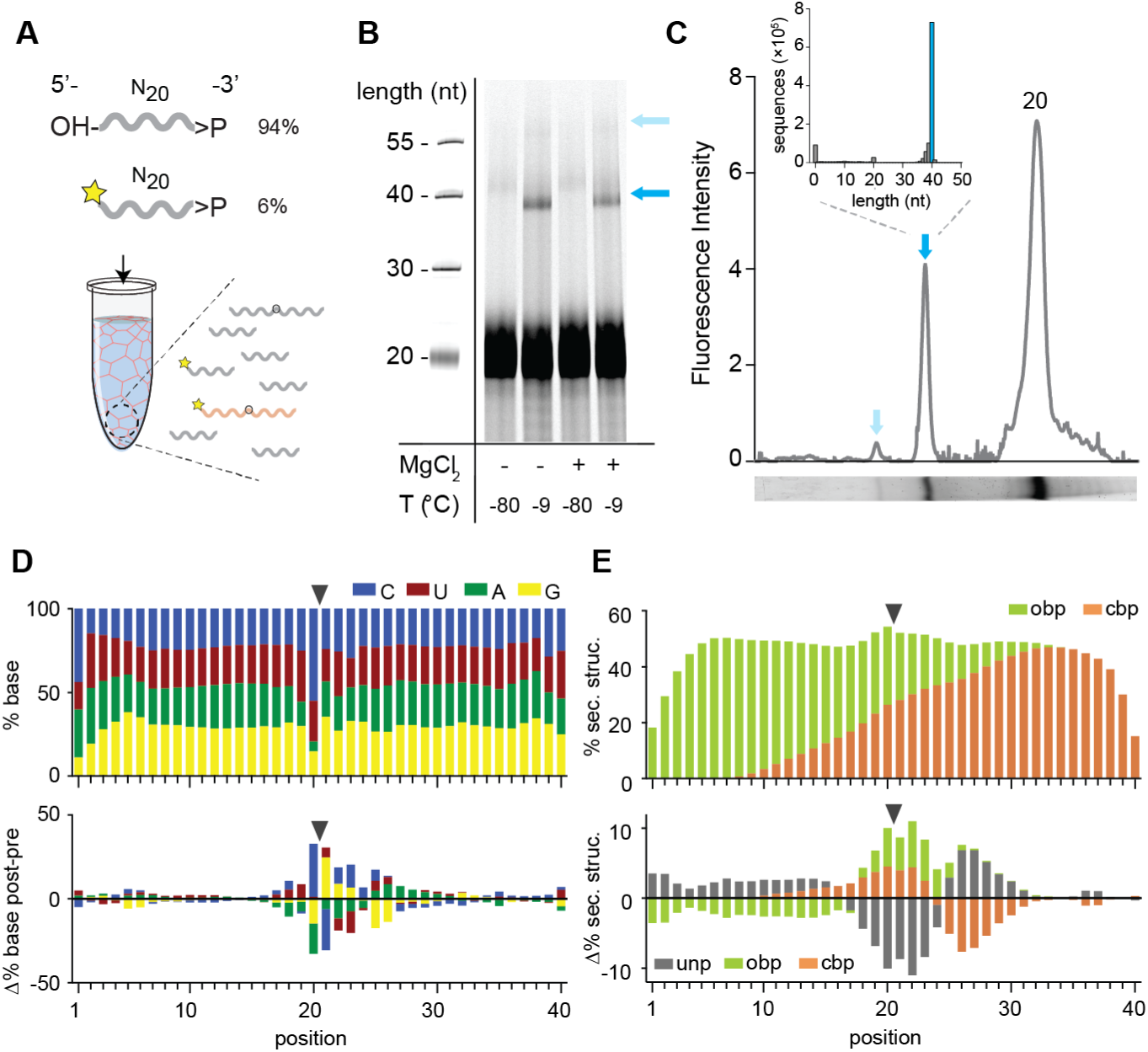
Reactivity of N20>p RNA pools. **(A)** N_20_>p pools are incubated in eutectic ice phases before analysis. 6% 5’FAM-labeled N_20_>p is spiked into the reaction to facilitate detection of ligation products (orange) by Urea-PAGE. **(B)** Scan of a denaturing gel for FAM-labelled products after 57 days of incubation either in eutectic ice (-9 °C) or at -80 °C, and +/- MgCl_2_. Note that ligation of FITC-N_20_>p is decreased due to the blocked 5’-OH. **(C)** Densitogram of a SYBR-gold stained reaction (-MgCl_2_, -9 °C) separated by Urea-PAGE monitoring also ligation of unlabeled strands. Arrows indicate ligation products. The size distribution of the main ligation product from deep sequencing implies direct ligation of two eicosamers (inset). **(D)** Upper panel: Average nucleotide distribution profile of the 40mer ligation products. Lower panel: Changes in nucleotide composition compared to the unligated input N_20_>p. Black arrows indicate the ligation site. **(E)** Upper panel: Average base-pairing frequencies of all sequenced 40mers as predicted by RNAfold (*10*) with opening base pairs in green and closing base pairs in orange. Lower panel: Difference of predicted base pairing between experimental and synthetic sequences generated *in silico* using the experimental nucleotide frequencies. Grey indicates predicted unpaired bases.

Interestingly, comparison of the 40mer ligation products with pre-ligation eicosamers revealed a clear nucleotide bias at the N_-1_pN_+1_ ligation junction (CpN) (Fig. 1D, Fig. S1A). Although mechanistically obscure, this sequence signature allows assignment of ligation junctions and reaction trajectories proceeding through a putative >p intermediate in diverse RNA contexts (see later). *In silico* RNA folding (*16*) of the ligated vs. unligated sequence pool indicated an increased intermolecular base-pairing tendency with ca. 55% of the intermolecular ligation junctions within helical regions (Fig. 1E). This suggests that ligation is accelerated by inter-strand hybridization-induced proximity, which both increases the effective concentration of reactive groups and reduces the entropic cost of the ligation reaction. Thus random pools provide organizing templates for ligation by intermolecular self-organization (*12, 13*).

We next sought to evaluate RNA pools for motifs that promote ligation by mechanisms other than simple intermolecular duplex formation. To reduce complexity and facilitate motif discovery, we repeated the ligation experiment using semi-randomer (srRNA) pools comprising random eicosamers flanked by (unstructured) constant sequence 20 nt primer binding sites (C1, C2). The use of “bait” 5’-C1-N_20_>p and “prey” 5’OH-N_20_-C2 oligonucleotides (2.5 µM each) enabled direct and rapid scoring of intermolecular ligation by RT-PCR using primers C1 and C2 after much shorter in-ice incubation times (Fig 2A, C; Fig. S2). Deep sequencing revealed ligation of srRNA 40-mer pools into 80-mer ligation products, with similar features as observed for the fully random pool (Fig. S1, S2), such as the characteristic ligation junction “fingerprint” and base-pairing bias. Applying a more stratified analysis, we searched for alternative RNA motifs at the ligation junction that may have mediated RNA ligation by comparing ligated bait-prey sequences from two different srRNA pools (differing in their C1, C2 sequences, Table S1). The use of different C1 and C2 anchor sequences allowed us to dissociate motif structure from sequence-dependent effects. We clustered ligation junction structures according to (predicted) minimal free energy using 408 different secondary structure sub-motifs based on 25 main-motifs including internal loops and hairpins (Fig. S3). While ligation frequencies of most of these motifs correlated with predicted base pairing frequency at the ligation junction (consistent with proximity effects), several submotifs showed high ligation frequencies inside internal loops (Fig. S4A, B). After further analysis using the motif discovery tool DREME (*17*), we chose two RNA motifs (H4 & J4) for further investigation, both comprising internal loops with well-defined consensus sequences (Fig. S4C, E). Both motifs were sufficient to promote ligation of minimal hairpin constructs (Fig. S4C - F): A minimized version of the J4 motif promotes reversible and regiospecific ligation via 2’-5’ regiochemistry both in ice (in absence of Mg^2+^) and ambient conditions (strictly Mg^2+^-dependent) with a ca. 10-fold enhanced apparent ligation rate (k_obs_) compared to a simple splint-mediated reaction (Fig. S5, S6) under the latter conditions. J4 is predicted to form a purine-rich 4×4 internal loop with a triple-sheared GA motif, which is notably also found in some natural RNAs (Fig S7). By contrast, the H4-motif mediates ligation via canonical 3’-5’ regiochemistry exclusively under eutectic ice conditions in the absence of Mg^2+^ but with a 2 to 4-fold slower k_obs_ than the splint-mediated (2’-5’) reaction (Fig. S5, S6).

**Fig 2.**
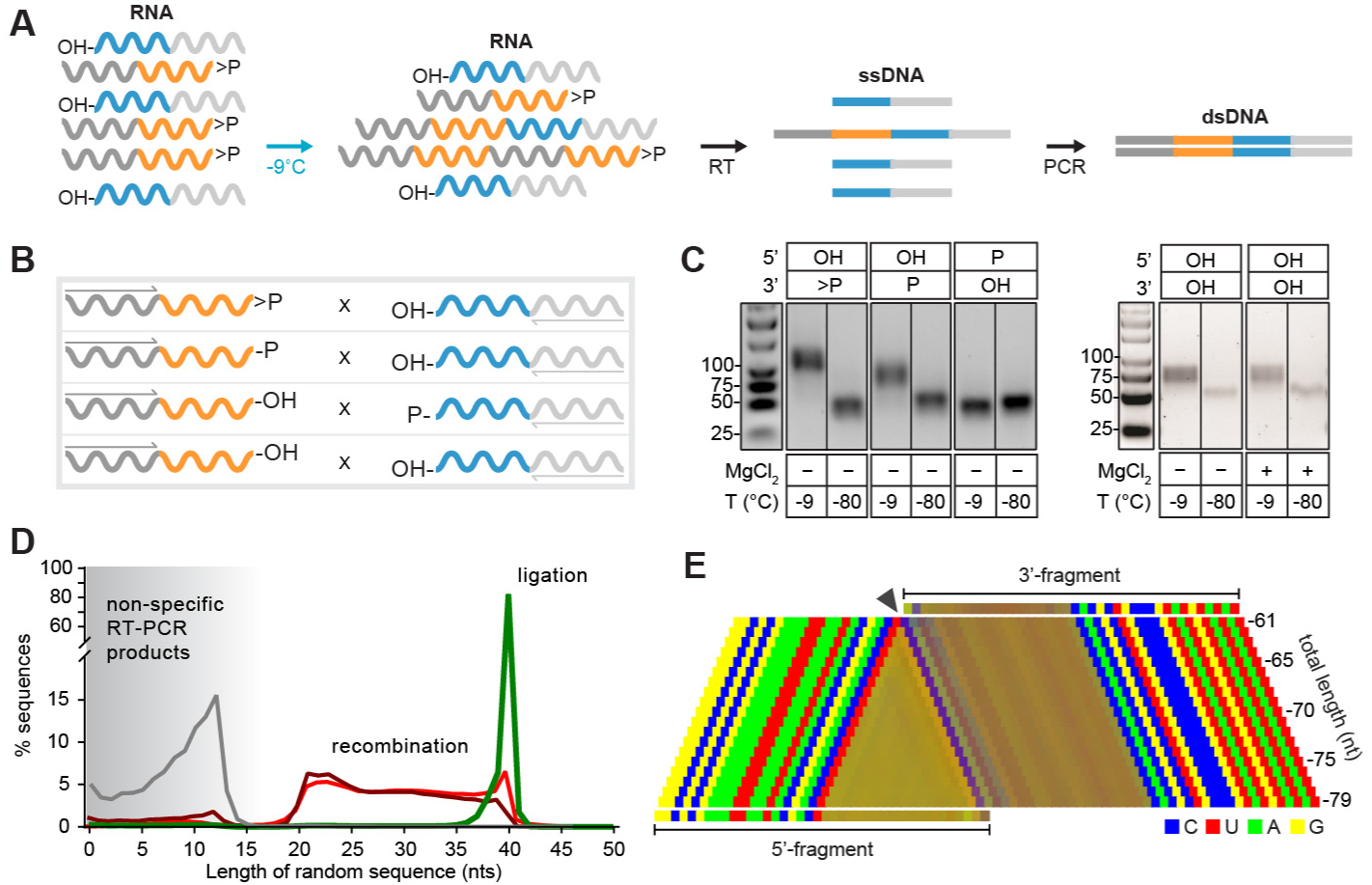
Reactivity of semi-random (sr)RNA pools with different 3’ and 5’ termini. **(A)** Assembly scheme: The random regions of both srRNAs are illustrated in orange (bait) and blue (prey). Only direct ligation products comprising 5’C1 (dark grey) and 3’C2 regions (light grey) are detected by RT-PCR. **(B)** Different srRNA bait / prey combinations. **(C)** RT-PCR results after incubation at -9 °C or -80 °C. Products recovered at -80 °C result from unspecific annealing of the RT-primer in the 3’-random region of the bait fragment (Fig. S2). **(D)** Size distribution of the RT-PCR products resulting from different srRNA bait × prey combinations. Combinations are C1-prey>p × 5’OH-bait-C2 (green), C1-bait-2’/3’p × 5’OH-prey-C2 (dark red), C1-prey-OH × 5’p-bait-C2 (grey) and naïve srRNA pools C1-bait-OH × 5’OH-bait-C2 (red) **(D)** Nucleotide signature for srRNA products created by recombination of C1-prey-2’/3’P × 5’OH-bait-C2. The frequency of the four nucleotides is represented by a linear combination of RGB values (C-blue, U-red, A-green, G-yellow) resulting in mixed colors in random segments between the primer binding sites used for recovery (see Supplementary material). The characteristic CpN signature at the presumed recombination site (arrow) indicates that the 5’-end of the full-length bait-C2 ligates to different length fragments of C1-prey.

Next, we explored mechanistic modes of intermolecular RNA ligation. Binary combinations of srRNA bait × prey pools (2.5 µM each) with different 2’, 3’- or 5’-substituents for ligation were compared, comprising C1-bait>p, C1-bait-2’/3’-monophosphate (C1-bait-2’/3’p), prey-C2 5’-monophosphate (5’p-prey-C2) as well as naïve srRNA pools (C1-bait-3’OH, 5’OH-prey-C2) (Fig. 2B). As expected, we observed ligation of C1-bait>p × 5’OH-C2-prey pools (and no ligation in negative controls incubated at -80 °C), as judged by RT-PCR (Fig. 2C). Furthermore, we observed intermolecular ligation in bait-2’/3’p × 5’OH-prey reactions, although these predominantly yielded shorter products, varying between 40 - 80 nts in length (Fig. 2C, D). These displayed the characteristic ligation “fingerprint” suggestive of reaction via a >p intermediate (see above), allowing mapping of ligation junctions. This revealed that variable-length assembly products derived from progressive truncation of bait segments. In contrast, prey segment length was preserved (Fig. 2E, Movie S1), suggesting intermolecular recombination (rather than ligation) via a transesterification mechanism presumably involving a nucleophilic attack of the prey 5’ OH on bait segments truncated by hydrolysis (Fig. S8). Consistent with this mechanistic hypothesis, blocking the prey pool 5’OH by phosphorylation (5’p-prey-C2, Fig. 2B) abolished detectable recombination (Fig. 2C). Furthermore, qRT-PCR suggests that recombination proceeds with a >10-fold slower rate than direct >p-dependent ligation (Fig. S9), consistent with a multi-step reaction trajectory requiring initial generation of an activated bait>p intermediate. Unexpectedly (but consistent with the above), even naïve srRNA pools (C1-bait-3’OH × 5’OH-prey-C2) – i.e. devoid of phosphorylation or activation chemistry - showed intermolecular assembly, yielding a similar distribution of truncated products that again included the characteristic junction fingerprint suggestive of a >p recombination intermediate.

In order to better understand naïve RNA pool reactivity without the imprint of conserved sequences, we next investigated naïve random RNA eicasomer pools, with only the prey including an invariant A_10_ tail sequence to facilitate recovery and analysis (bait × prey: 5’FAM-N_20_-3’OH × 5’OH-N_20_A_10_-3’OH). After an extended incubation (5 months, -9 °C) we were able to directly observe formation of two distinct recombination products by gel electrophoresis without any amplification (Fig 3A). The two main product bands (P1 & P2, together ca. 2% yield) were analyzed using SMARTer™ RT-PCR and deep sequencing (Fig S10), revealing recombination fragment distributions from ~40 to ~50 nts (for P1) that include the characteristic fingerprint of a >p ligation junction (Fig 3B, C). In contrast, the longer (on average) P2 products (~45 to ~55 nts) showed an ApN ligation junction as they were formed (albeit less efficiently) from recombination of two prey segments within the A_10_ tail (Fig 3B, C). Thus, as with naïve srRNA pools, intermolecular ligation products are likely formed by a two-step transesterification process initiated by hydrolytic cleavage (of either bait or prey), followed by putative ligation of shortened >p sub-fragments, giving rise to distinct product species such as bait-prey (P1) or prey-prey (P1 and P2). Bait-bait recombination is not observed, presumably as the bait 5’ OH is FAM modified to aid analysis. Comparison of P1 / P2 pools with random pools of equal nucleotide distribution generated *in silico* also suggested that recombination sites were preferentially localized in base-paired regions (Fig 3D) and that selected pools exhibited an overall increase in (predicted) folding propensity and stability (Fig 3E). Thus, naïve random-sequence RNA pools, devoid of any activation chemistry or conserved sequence regions (apart from the prey A_10_ tag) have an innate capacity to self-organize and spontaneously recombine yielding subpools with increased oligomer length, sequence diversity and propensity to form secondary structures. This phenomenon may be a crucial factor in the emergence of longer molecules with *inter alia* distinct ribozyme activities from prebiotic pools.

**Fig. 3.**
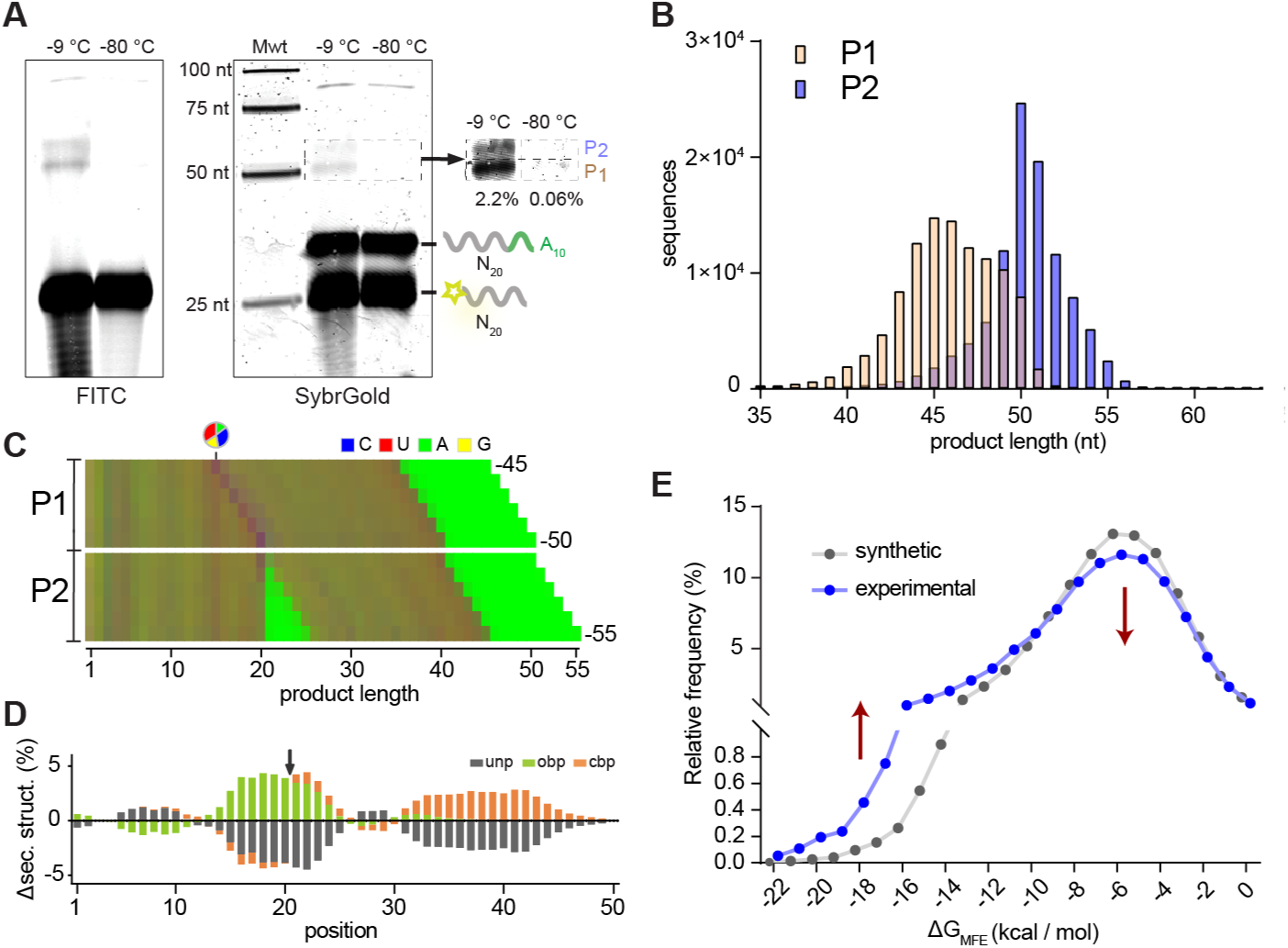
Reactivity of naïve random RNA pools. **(A)** Denaturing gel electrophoresis of naïve N_20_A_10_ and FAM-N_20_ pools after incubation (-9 °C or -80 °C, 5 months). Products lengths show two main product bands (P1, P2). **(B)** Size distributions of the sequenced P1 (yellow) and P2 (blue) band. **(C)** Color-coded nucleotide frequencies of P1 and P2 sequences recovered using the protocol shown in (Fig. S10). **(D)** Average base-pairing differences between the experimental 40mer product pool and randomly generated *in silico* sequences using the experimental nucleotide frequencies (C). Secondary structures were calculated using RNAfold. Grey indicates predicted unpaired (unp) nucleotides, green nucleotides involved in opening base pairs (obp) and orange closing base pairs (cbp). **(E)** Theoretical energies of MFE structures from experimental and *in silico* pools also used in (D) suggest that recombination product pools show enhanced folding stabilities.

Finally, we sought to dissect chemical parameters critical for an innate recombination potential. To this end, we expanded our investigations beyond RNA to compare and contrast the potential of random-sequence pools of a range of RNA congeners (representing systematic variations of the canonical ribofuranose ring structure, also known as xeno nucleic acids (XNAs)) with respect to intermolecular ligation or recombination. We first examined naïve semi-randomer pools of DNA or ANA (arabino nucleic acid) (*18*), i.e. replacing the canonical ribofuranose ring of RNA with either 2’-deoxyribose (DNA) or arabinofuranose (ANA), in which the 2’ hydroxyl group of RNA is in the axial (trans) position. Like RNA and DNA, ANA is capable of forming functional catalytic motifs (*19*). Furthermore, several small self-cleaving DNA sequence motifs have been identified (*20*), suggesting that, for DNA at least, there should be a number of pool sequences able to perform the first step of a recombination reaction. However, while we readily detected recombination in the naïve srRNA pools (as above), neither srDNA nor srANA pools showed detectable reactivity even after prolonged incubation (up to 6 months, -9 °C; Fig. 4A) and despite the 10-100 higher sensitivity of PCR (DNA) compared to RT-PCR (RNA & ANA) detection (*21, 22*). Likewise, inclusion of transition metal ions (Zn^2+^), which have been shown to promote DNA self-cleavage (*20*) did not result in recombination in DNA (or ANA) pools, while apparently inhibiting it in RNA pools (Fig S11).

**Figure 4.**
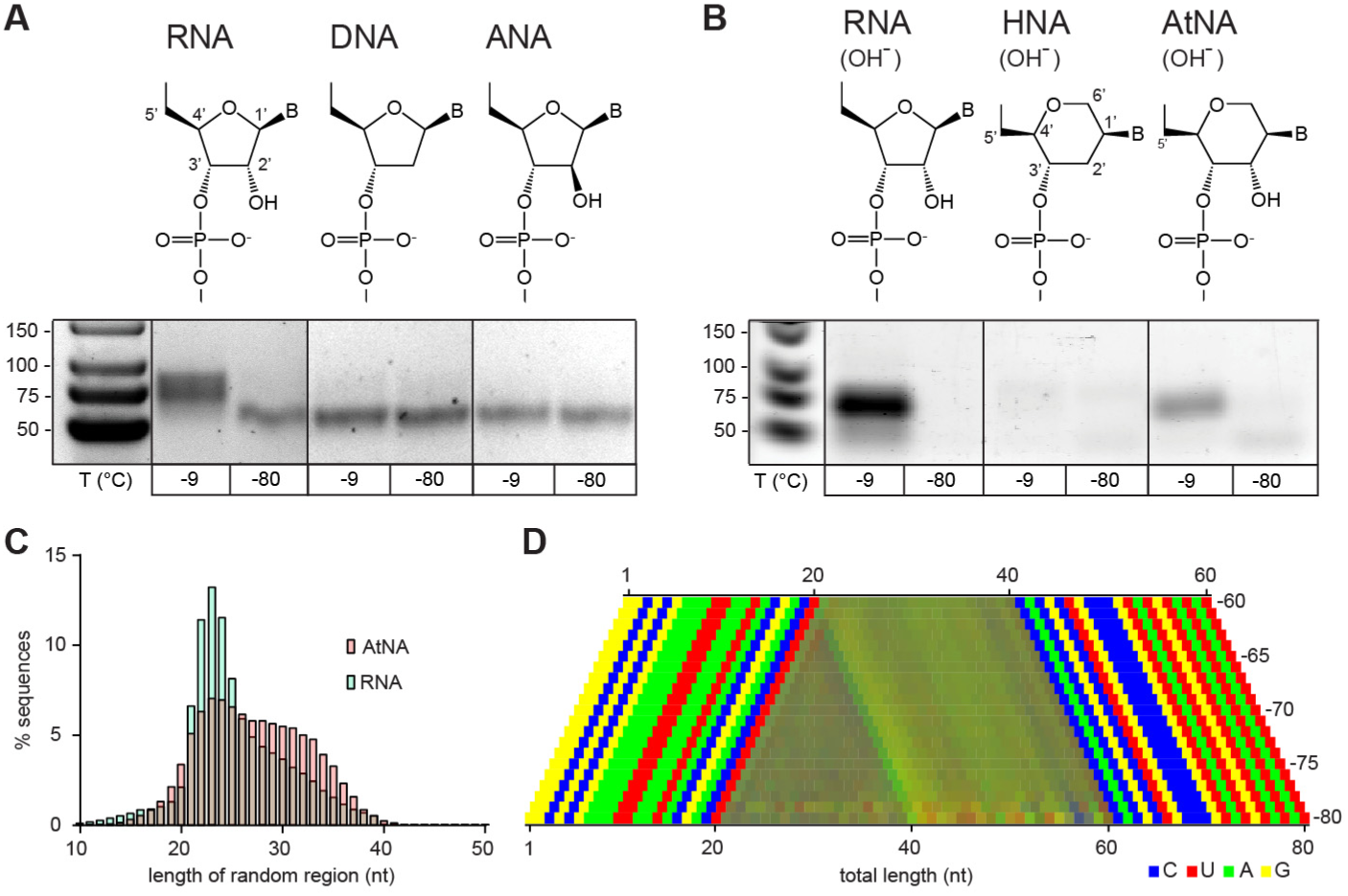
Reactivity of naïve semi-random pools of genetic polymers. **(A)** In contrast to srRNA, srDNA and srANA pools show no spontaneous recombination (-9 °C, 5 months). **(B)** Naïve srRNA and srAtNA pools recombine, while HNA shows no recombination (after a brief NaOH treatment). **(C)** Size distributions of the srRNA and srAtNA products. srRNA pools are biased towards shorter fragments, presumable due to hydrolysis during NaOH treatment. **(D)** Color-coded nucleotide signatures of srAtNA products from (B) shows similar bait segment truncation as in RNA (see Fig. 2) but with a distinct ApN ligation signature.

We next explored the reactivity of pools of two additional RNA congener XNAs: D-altritol nucleic acid (AtNA), which shares an analogous vicinal 2’, 3’-cis-diol configuration with RNA but in a six-membered ring (Fig 4B), as well as its 2’-deoxy-analogue, hexitol nucleic acid (HNA). To test whether the AtNA vicinal hydroxyl can participate in transesterification reactions despite its different ring geometry, we exposed defined-sequence HNA and AtNA oligonucleotides as well as equivalent RNA to NaOH. We found that AtNA (like RNA) is sensitive to alkaline pH-induced hydrolysis, albeit with an apparent 6-fold slower strand scission rate than RNA, while HNA proved inert (Fig S12). This indicates that (akin to RNA) the equivalent vicinal 2’OH in AtNA can participate in intramolecular transesterification reactions presumably via an analogue of the 2’, 3’-cyclic phosphate (AtNA>p) intermediate of RNA (Fig S8C); the observed lower hydrolytic reactivity suggests that formation of the AtNA>p intermediate may be energetically less favorable.

Finally, we sought to test whether semi-randomer pools of HNA (srHNA) and AtNA (srAtNA) could participate in intermolecular recombination reactions. To this end, we compared the reactivity of srHNA and srAtNA pools to equivalent srRNA pools. To accelerate reactivity, we exposed srHNA and srAtNA (as well as control srRNA) pools to a short pulse of 50 mM NaOH at 65 °C (RNA (3 min), HNA / AtNA (27 min), neutralized with 1 M Tris∙HCl pH 7.4) before incubation in ice for 5 months. Using a novel engineered reverse transcriptase (RT TK^2^, see Methods), which (unlike commercial enzymes) is able to reverse transcribe all three XNA chemistries, we analyzed ligation products by RT-PCR. Clear intermolecular ligation and recombination was again observed in the srRNA pool, but also in the srAtNA pools, with a similar size distribution but distinct nucleotide signature (Fig 4B-D). In contrast, srHNA pools proved inert. These findings suggest that a vicinal cis-diol configuration may be a critical determinant of an innate intermolecular recombination potential regardless of ring geometry.

## Discussion

Repertoire selection experiments have previously shown that random sequence pools of different genetic polymers (including RNA, DNA and some XNAs) comprise rare, functional sequences capable of strand cleavage and / or ligation, which can be isolated by iterative rounds of selection and amplification (*23*). Here, we show that in some genetic polymers, the pools themselves show an innate, global potential for spontaneous recombination. Although to some degree this involves specific sequence motifs that (e.g. in RNA pools) promote ligation with either 2’-5’ or 3’-5’ regioselectivity (Fig S4 - S5), the bulk of observed reactivity instead derives from molecular self-organization of pool oligomers into intermolecular non-covalent complexes. In the case of pools of random RNA as well as semi-random RNA and AtNA oligomers, a subset of these intermolecular assemblies do not remain inert (and non-covalent), but have the capacity to undergo spontaneous ligation and recombination reactions through transesterification chemistry as detected by gel shift in denaturing gel electrophoresis and specific RT-PCR amplification.

While we observe efficient intermolecular ligation and recombination of RNA after just a few days in-ice incubation, we note that the simple band-shift and RT-PCR assay we use likely underestimates total pool reactivity, as a range of more complex recombination modes, including intra- and intermolecular circular, lariat or branched products are likely to be missed (Fig S13). At the same time, our assays provide a subset of *bona fide* true ligation and recombination products as PCR or as sequencing artefacts can be ruled out on multiple grounds. For one, incubation at -80 °C, which freezes the eutectic phase in our buffer system (*7, 24*), never yields products, establishing that fragments do not recombine during workup. Furthermore, the appearance (or absence) of recombination products is clearly and reproducibly linked to chemical features of the original pool (e.g. free 5’OH (RNA), or cis diol (RNA, AtNA)), which are erased in the RT step. Finally, in both the case of the >p-activated and naïve RNA randomer pools, we are able to observe the formation of recombination products directly by denaturing gel electrophoresis, which would dissociate even highly stable non-covalent complexes (Fig 1, 3).

Previous reports of non-enzymatic RNA recombination with defined substrates suggests that the majority of such reactions are likely to involve a two step-mechanism of two consecutive transesterification reactions via an >p intermediate. These may be promoted by proximity through *in trans* hybridization and the concentration and entropic effects of eutectic ice phase formation (*25–27*) (Fig. S8). Whilst formation of canonical 3’-5’ linkages was observed with some motifs (H4), we expect the majority of RNA pool linkages to conform to 2’-5’ regiochemistry as such linkages form preferentially in RNA duplex geometry (*28*), although such information is lacking for AtNA. In RNA, sporadic 2’-5’ linkages have been found to be broadly compatible with folding and function (*29*) and can progressively be converted to the canonical 3’-5’ linkage (*30*).

The mechanistic basis for the distinctive sequence signature (CpN) in RNA is unclear. CpA and UpA junctions have been observed to be more reactive (*31*), thus the bias may reflect the higher propensity for C>p formation, but this can be confounded by dramatically altered reactivity depending on local sequence and chemical (H-bonding) context (*31, 32*), which may explain the different preferred signature (ApN) in the case of AtNA (Fig. 4D).

We observe striking differences in innate reactive potential of the different genetic polymers RNA, DNA, ANA, HNA and AtNA. The previously reported isolation of highly active DNAzymes (*33*) and to a lesser extent ANA-, HNA- and other XNAzymes (*19*) by *in vitro* evolution clearly indicate that such pools do contain functional sequences. This raises the question of why DNA, ANA and HNA pools all proved inert in our recombination experiments even in the presence of transition metal ions like Zn^2+^, which had been shown to promote self-cleavage in DNA (*20*). While nucleophilicity and p*K*_a_ of the 5’-OH are likely to be comparable in these different genetic polymers, studies using model nucleotide phosphodiester and -triester compounds suggest an at least 30-fold higher reactivity of RNA internucleotide diester linkages compared to DNA and ANA (*34*). The latter is thought to be due to the vicinal cis diol configuration of RNA, with the 2’-OH stabilizing the negative charge developing on the phosphorane intermediate and / or the departing 3’-oxygen by H-bonding (*32, 34*) or local electronic effects (*35*), with reactivity furthermore highly dependent on local sequence context, stacking and basepairing effects (*31*). Consistent with this, the RNA congener AtNA, sharing a vicinal cis-diol configuration (in a six membered ring context), was found to be capable of analogous intermolecular ligation and recombination, although AtNA reactivity appears to be significantly lower than RNA. The latter may due to greater torsional strain required for in-line attack in formation of an analogous 2’, 3’-cyclic phosphate (AtNA>p) intermediate (with an almost planar O2-C2-C3-O3 configuration) in the context of a six-membered ring (*36*).

Although >p is only weakly activated towards nucleophilic attack, in the context of high effective concentrations of 5’ OH nucleophile in a gapped duplex (or pre-organized internal loop motifs like H4, J4 (Fig. S4) or bulges), bond formation can be efficient. Crucially, >p groups are readily formed as part of RNA (and presumably AtNA) hydrolysis and potentially as products of both *in cis* and *in trans* nucleophilic attack of RNA 5’-, 2’- and 3’-OH groups leading to the observed recombination reactions. While DNA pools comprise a number of small self-cleaving motifs (*20*), these yield 3’OH and 5’p termini, which are not competent for ligation without further activation e.g. with a phosphorimidazoide (*37*). Thus a vicinal diol configuration not only accelerates the initial strand cleavage step of recombination but also yields cleavage product termini (>p and 5’OH) competent for re-ligation. Other genetic polymers comprising vicinal cis diols include the ribo- and lyxopyranosyl series (*38, 39*), which may therefore be similarly predisposed to recombination (Fig S14). Indeed, ligative polymerization of 2’, 3’ >p activated pyranosyl tetranucleotides on a template has demonstrated (*40*).

In conclusion, we observe that naïve random and semi-random RNA (and AtNA) oligomer pools undergo spontaneous recombination among pool sequences, generating a distribution of new recombined oligomers. Such recombination reactions progressively increase pool complexity by the *de novo* generation of diversity in oligomer length, sequence content and secondary structure. This is significant in the context of previous arguments on the importance of oligomer length for the encoding of the most active larger, functional motifs (*2, 41, 42*) (or more frequent encoding of smaller motifs), as well as the importance of sequence diversity and recombination in accelerating adaptive walks (*43–46, 7*). These findings suggest that such naïve random RNA and AtNA pools are predisposed to “bootstrap” themselves towards higher compositional and structural complexity (*47*) and thus a likely enhanced adaptive potential.

## Acknowledgments

The authors thank J. Attwater for discussions and comments on the manuscript. **Funding:** This work was supported by a Federation of European Biochemical Societies (FEBS) Long-Term Fellowship (HM), the Medical Research Council (PhH, AIT, GH, program no. MC_U105178804) and from KULeuven, Research Council (OT/1414/128) and FWO Vlaanderen (G078014N) (MA and PiH). **Author contributions:** HM, AIT and PhH conceived and designed the experiments, analyzed data and wrote the manuscript. AL contributed to H4 and J4 motif identification and their characterization. GH contributed the new RT-enzyme, which will be described in detail elsewhere. MA and PiH synthesized the HNA and AtNA nucleosides. **Competing interests:** Authors declare no competing interests. **Data and materials availability:** Data is available in the main text or the supplementary materials. Sequencing files and analysis scripts are available upon request.

## Supplementary Materials

Materials and Methods

Figures S1-S13

Tables S1-S3

Movie S1

References (*7, 16, 17, 19, 24, 48–60*)

